# The most remarkable migrants – systematic analysis of the Western European insect flyway at a Pyrenean mountain pass

**DOI:** 10.1101/2023.07.17.549321

**Authors:** Will L Hawkes, Toby Doyle, Richard Massy, Scarlett Weston, Kelsey Davies, Elliott Cornelius, Connor Collier, Jason W. Chapman, Don R. Reynolds, Karl Wotton

## Abstract

In 1950 David and Elizabeth Lack chanced upon a huge migration of insects and birds flying through the Pyrenean Pass of Bujaruelo, later describing the spectacle as combining both grandeur with novelty. The intervening years have seen many changes to land use and climate, posing the question as to the current status of this migratory phenomenon, while a lack of quantitative data has prevented insights into the ecological impact of this mass insect migration and into the factors affecting it. To address this, we revisited the site in autumn over a 4-year period and systematically monitored diurnal insect species and numbers. We document an annual mean of 17.1 million day-flying insects from 5 orders moving south, with ‘mass migration’ events associated with warmer temperatures, the presence of a headwind, sunlight, low windspeed, and low rainfall. Diptera dominated the migratory assemblage and annual numbers varied by more than fourfold with larger annual migration flows associated with higher autumn temperatures in Northwest Europe. Finally, using observed environmental thresholds for migration, we estimate an annual ‘bioflow’ of at least 14.6 billion day-flying insects migrating south over the whole Pyrenean Mountain range, highlighting the importance of this route for seasonal insect migrants.

## Introduction

Insects are the most numerous terrestrial migrants, far surpassing vertebrates in terms of biomass and abundance (1,2). Insects migrate to exploit seasonal resources, increase reproductive output, and/or escape habitat deterioration, e.g., due to temperature change, food quality or disease risk, or to seek sites to overwinter (1,3). The life histories of North Temperate Zone migrants are often associated with a high reproductive advantage such that autumn migrants considerably outnumber spring influxes to northern latitudes, for example by an average of over twofold for hoverflies and over threefold for *Autographa gamma* (silver Y moth) (4,5).

Insect migration studies are often limited to specific taxa, namely Lepidoptera, dragonflies and hoverflies (6–8), while whole assemblage studies are relatively rare. This is because small insect migrants are often invisible in a large landscape necessitating specialist equipment such as entomological radars (9,10) or balloon-supported nets for their detection (2,11,12). However, in some rare locations known as migration hotspots (also referred to as bottlenecks) aerial densities are greatly increased by topographical conditions, allowing the whole migratory assemblage to be studied at ground level. Relatively few insect migration hotspots are known, but where studies have been made valuable information about types and numbers of migrating insects has been obtained (8,13–15). In Europe during the autumn, migration hotspots can be found within mountain ranges such as the Alps in Central Europe and the Pyrenees in Western Europe (8,15); in these locations, mountain barriers and local winds direct insects up steep-sided valleys and through narrow mountain passes. In Central European migration hotspots, insects have been monitored at Randecker Maar research station in the Schwäbische Alb uplands in Germany since 1978, and over the Col de Bretolet mountain pass in the Swiss-French Alps for over 12 years between 1962 to 1973 (8,16).

In contrast, Western European migration hotspots have been poorly studied for insects despite the early recognition of these flyways (13,15,17,18). The most significant of these passes is Puerto de Bujaruelo (Port de Gavarnie) in the Haute-Pyrenees, where David and Elizabeth Lack ‘chanced upon a spectacle’ of insect migration on a single day in mid-October 1950 and recorded their observations (15). This classic study revealed movements of butterflies, dragonflies, and huge numbers of hoverflies, describing these as ‘the most remarkable migrants of all’ due to their sheer abundance. Between 1951 and 1952, several researchers briefly visited the pass, and elsewhere in the Pyrenees, and noted evidence of migration from Lepidoptera, Diptera, and Odonata (17–19). In 1953, Williams et al (13) studied insect migration at three sites in the Pyrenees including the Bujaruelo Pass from 18 September to 15 October, making it the longest continuous observational study of insect migrants through a Western European migration hotspot. They confirmed previous observations and added Hymenoptera to the list of migratory taxa, as well as recording *ad lib.* meteorological conditions leading to the conclusions that insects do not migrate on cold, wet, and sunless days (13). While these studies made important contributions to the field, they lacked a systematic and sustained approach to the recording of insect migration, leaving the question as to the diversity of migrants using this route, the number and variation in individuals, and how this variation is affected by environmental factors. To rectify this gap in our knowledge, we systematically quantified daytime insect migration through the Bujaruelo Pass during the September to October migration period in the years 2018 to 2021.

## Materials and methods

### Location

The Pyrenees Mountain range runs the border between southern France and Northeast Spain (Figure 1A). Within the Hautes-Pyrénées region of southwest France, the Vallées des Gaves runs north to south from the town of Argeles-Gazost before ending abruptly at the high-walled Cirque du Gavarnie (Figure 1B). A smaller valley runs northeast to southwest from the Cirque du Gavarnie, culminating in the mountain pass of Bujaruelo where we carried out our observations (42°42’14.2“N 0°03’51.4”W) (Figure 1B). The pass itself sits at 2273 m and is 30 m wide hemmed in by high mountain peaks (Pic Entre les Ports – 2476 m and Pic de Gabiet – 2716 m to the north, and El Taillon – 3144 m to the south; Figure 1B). The northeast part of the pass falls within the Parc National des Pyrénées and the area consists predominately of rocky shale with patches of grass kept short through weather and grazing pressures. On the slopes leading up to each side of the pass, flowering plants grow with Pyrenean thistle (*Carduus carlinoides*) being the only nectar source in substantial numbers. All fieldwork occurred between the months of September and October annually during 2018 to 2021. Permission to conduct observations was obtained from the Parc national des Pyrénées (France, authorisation numbers: 2018-9, 2019-67, 2020-146, and 2021-33) and the Gobierno de Aragon (Spain, authorisation numbers: 500201/24/2018/06141, 500201/24/2019/02174, 500201/24/2020/01724, and 500201/24/2021/01722).

**Figure 1.**
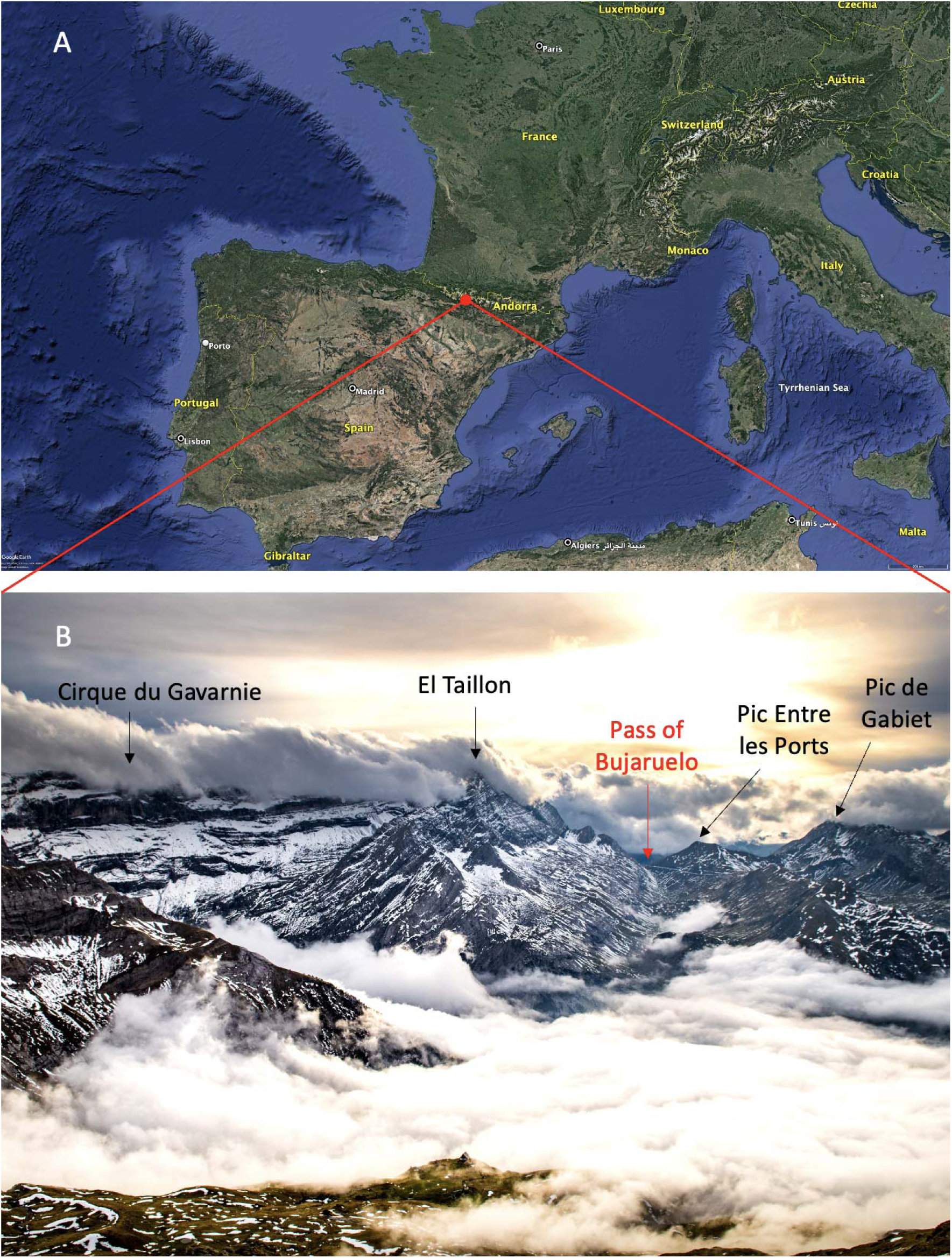
The Pyrenean Mountain range and the Pass of Bujaruelo. (A) The fieldwork location (red marker) within the Pyrenean Mountain range. (B) The valley which channels the insects over the Pass of Bujaruelo (red arrow) and the surrounding geological formations.

### Insect identification

Identification of migrating butterflies and dragonflies were carried out by eye as they flew over the pass. Smaller insects were caught with a bidirectional malaise trap continuously set up on the pass with openings to the southwest and northeast (ez-Migration Trap, Bugdorm). Samples from this trap were collected each evening and identified to at least family level under a Leica MZ6 dissecting microscope. During the study periods of 2020 and 2021, the sex ratios of three species (the hoverflies *Melanostoma mellinum* and *Eupeodes corollae*, and the muscid fly *Musca autumnalis*) were also recorded. Migrants were determined based on a ‘southward score’ which is the percentage caught heading south in a wind category (headwind or tailwind) minus the percentage caught heading north in the same wind category. For full details of this methodology, please see the Supplementary Methods.

### Quantifying the bioflow of the larger migrants

The movement of the migrant butterflies and dragonflies were quantified by counting numbers as they passed through the 30m wide pass during a 15-minute period, once every two hours from 10:00 to 16:00. These numbers were then scaled for the remaining two hours between surveys up until 18:00 by multiplying the counts of insects (N1) by eight to reach two hours.

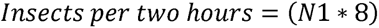

The migratory traffic rate (MTR: individuals per metre per minute) of the insects was calculated by dividing the number of insects recorded in a count (N1) by the number of minutes recorded (15 = min), before dividing this figure by the width of the pass (30m = W).

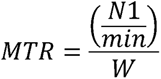

### Quantifying the bioflow of smaller migrants

Numbers of the smaller migrants were quantified from video recordings over a 2-m wide section of the pass as in Hawkes et al (14). We scaled these raw insect counts to get a more representative estimation of the insects migrating through the Bujaruelo pass (R1). To scale for time, raw counts (N2) were multiplied by 15 to obtain representative counts for 15 minutes. To scale for the width of the pass (30m) the counts were multiplied by 15 to encompass the entire width.

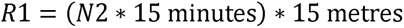

The migratory traffic rate (MTR) of the smaller insects was calculated by dividing the number of insects recorded in a count (N2) by the number of minutes recorded (1 = min), before dividing this figure by the width over which the counts was performed (2m = W).

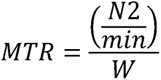

The video trap was unable to resolve insects smaller than 8-10 mm in length as was noted during previous research (14). Of the insects caught in the intercept traps that showed migratory behaviour, 57% were too small to be resolved on the video camera. To account for this, all mentions of insect counts below include the missed insects.

### Diurnal insect timings

To determine the timings at which insects fly during the day, 22 separate days from the 2019, 2020, and 2021 field seasons were chosen to analyse. These days contained ‘mass migration’ events. The ‘mass migration’ days were classified by ordering counts from all days across four years from highest to lowest and selecting as ‘mass migration’ events until they cumulatively reached 95% of the total number of individual insects recorded. The mean counts from every 15 minutes during these days (from 09:00 – 17:00) was then calculated to identify the daily temporal distributions of migrating insects.

### Meteorological data

Weather conditions were recorded every two hours between 10:00 and 16:00 each day. Measurements included cloud cover recorded in the Okta scale (UKMO, 2010), windspeed and wind direction in m/s using a Windmate Anemometer, percentage of unobscured sunlight, temperature in degree Celsius and precipitation as dry or raining. In addition, continuous weather data was obtained for the Pass of Bujaruelo from meteoblue simulations (20). Our in-situ meteorological recordings closely matched the meteoblue data and so this was used for analysis due to its continuous nature (20). Large scale wind patterns were visualised using Ventusky.

### Analysis of meteorological factors

To test which environmental factors affected the numbers of insects migrating through the pass, a generalised linear model (GLM) was fitted with a log link function and Poisson family (21). Counts of the insects were the response variable and the explanatory variables were [1] temperature, [2] total daily sunlight, [3] average daily windspeed, [4] total daily precipitation. To determine the significance of the explanatory variables, the variable of interest was excluded before comparing the models with and without the variables using ANOVA. To investigate the effect of meteorological conditions on annual variation of insect numbers, the numbers of insects recorded in the Pyrenees were correlated using Pearson’s correlation with meteorological data from a location in northwest Europe: Great Britain (the Met Office, UK). Further analysis to gain a more representative picture was performed using a UK-based ten-year entomological radar dataset(5). All statistical procedures were performed in R version 4.2.2. (22).

### Fine temporal scale effect of sunlight on insect migration

Days of mass migration events were chosen for sunlight analyses as all the other conditions affecting insect migration were likely to be similar across these events. Sunlight analysis was performed at a fine temporal scale on the mountain pass using the video trap footage from the 2021 field season to check for presence/absence of sunlight. Each of 382 occasions were classified as either sun present or absent and the number of insects within each measurement occasion was recorded.

### Quantifying numbers of insects annually traversing the Pyrenees

To quantify the numbers of insects passing over the entirety of the Pyrenees each autumn we identified the location of the 1.2 km wide valley mouth (located at: 43°04’46.9”N 0°02’43.4”W) where the migratory assemblage moving through the Pass of Bujaruelo would first begin to be channelled up the valley towards the pass. We made the assumptions that the number of insects entering the valley is equal to the number moving through the 30m wide pass, and that the rate of insects migrating is constant across the entire Pyrenees. Based on these assumptions we calculate the flow over the 430 km length of the Pyrenees based on the numbers at the pass, for full methodology, see Supplementary Methods.

## Results

### Assemblage of insects migrating through the Pyrenees

The complete assemblage of all migratory insects consisted of 20 families from 5 orders (Supplementary Tables S1-S4): Diptera (89% of total migratory insect number), Hymenoptera (6%), Hemiptera (5%), Lepidoptera (<1%) and Odonata (<1%) (Figure 2A). From the southward scores, determined from patterns of movement under different wind conditions, four insect types were classed as strongly migratory: hoverflies (Syrphidae), red admiral butterflies (*Vanessa atalanta*), ‘blue’ butterflies (Lycaenidae), and blowflies (Calliphoridae) (Figure 2B & C, Supplementary Table S2); eight insect types were classed as migratory: ‘white’ butterflies (*Pieris* spp.), clouded yellow butterflies (*Colias croceus*), and the Diptera families Muscidae, Anthomyiidae, Phoriidae, Chloropidae, Mycetophilidae, and Sciaridae) (Figure 2B & C, Supplementary Table S3); and four groups were classified as showing weaker migratory behaviour: the Diptera families Lonchopteridae and Sphaeroceridae, solitary Hymenoptera, and aphids (Aphidoidea) (Figure 2B & C, Supplementary Table S4). Classification details of the different migratory groups can be found in the Supplementary Methods. Further species and groups like the painted lady butterflies (*Vanessa cardui*), hummingbird hawkmoths (*Macroglossum stellatarum*) and some dragonflies (*Aeshna mixta* and *Sympetrum striolatum*) were seen performing directed movements southwards over the pass, but their numbers did not exceed the 100 individual threshold to be included in further analyses. It is thought that the numbers recorded of dragonflies and hummingbird hawkmoths are likely an underestimation, but their darker colouration and fast flight made them difficult to count. The primary ecological roles (occasionally insects had two or more primary roles) of the migratory assemblage were identified from the literature and revealed that the most numerous were pollinators (87.5% of total insects counted from both video camera and butterfly counts), with other, sometimes overlapping roles including decomposers (33.6%), pest predators (22.2%), and pests (5%), with 100% of the total insect assemblage contributing to nutrient transfer (Supplementary Table S5).

**Figure 2.**
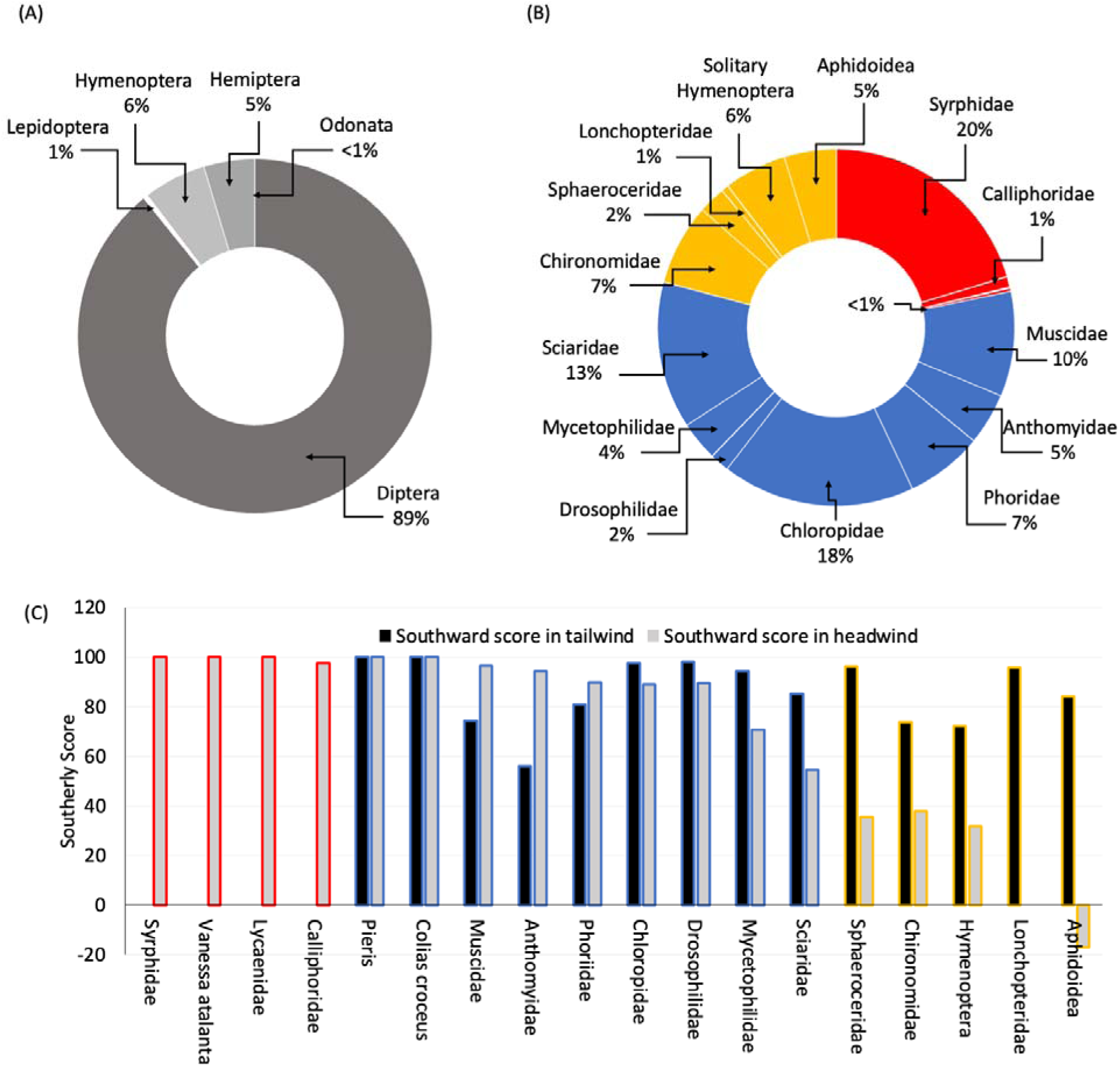
Classification of the migratory assemblage. Average ratios of insects showing migratory behaviour collected in the intercept trap and butterfly counts over 4 years sorted by (A) Order and (B) Family. (C) Migratory behaviour of insect groups based on southward scores. ‘Strongly migratory’ insect groups (red), ‘migratory’ insect groups (blue), ‘weakly migratory’ groups (yellow). Missing values indicate <100 individuals in the sample of head or tail winds. Southward scores of 100 indicate all individuals were going south, 0 that the same number were going south as north and negative vales indicate that more were going north than south under a particular wind condition.

### Quantifying insect bioflows through the Pass of Bujaruelo

In total, scaled counts from malaise traps, video traps, and butterfly counts reveal an average of over 17.1 million insects traverse the pass of Bujaruelo each autumn. However, variation from year-to-year ranged from 6.2 million insects to 27.1 million insects an annual variation of 4.4-fold (Figure 3A subset), full breakdown can be found in Supplementary table S6). The majority of these were the smaller insects recorded by the video camera which made up 99.6% of total migrants or 68 million individuals over the four years. The butterflies comprised only 0.4% of the total with just over 260,000 individuals across the four years with annual counts varying from 38,500 to 107,000. To allow a comparison between our data and other previously investigated migratory hotspots we calculated the migratory traffic rate given as the number of small insects per metre per minute. Numbers through the pass varied between 0 and 3683 individuals across the four years of data collection. Butterfly migratory traffic rates were much lower and varied between 0 and 4 individuals across the four years. The temporal distribution of all insects recorded by the camera trap over a day (09:00 -17:00) reveals the peak of insect numbers occurs at 14:45 (Figure S1), about 1 hour after the solar noon.

**Figure 3.**
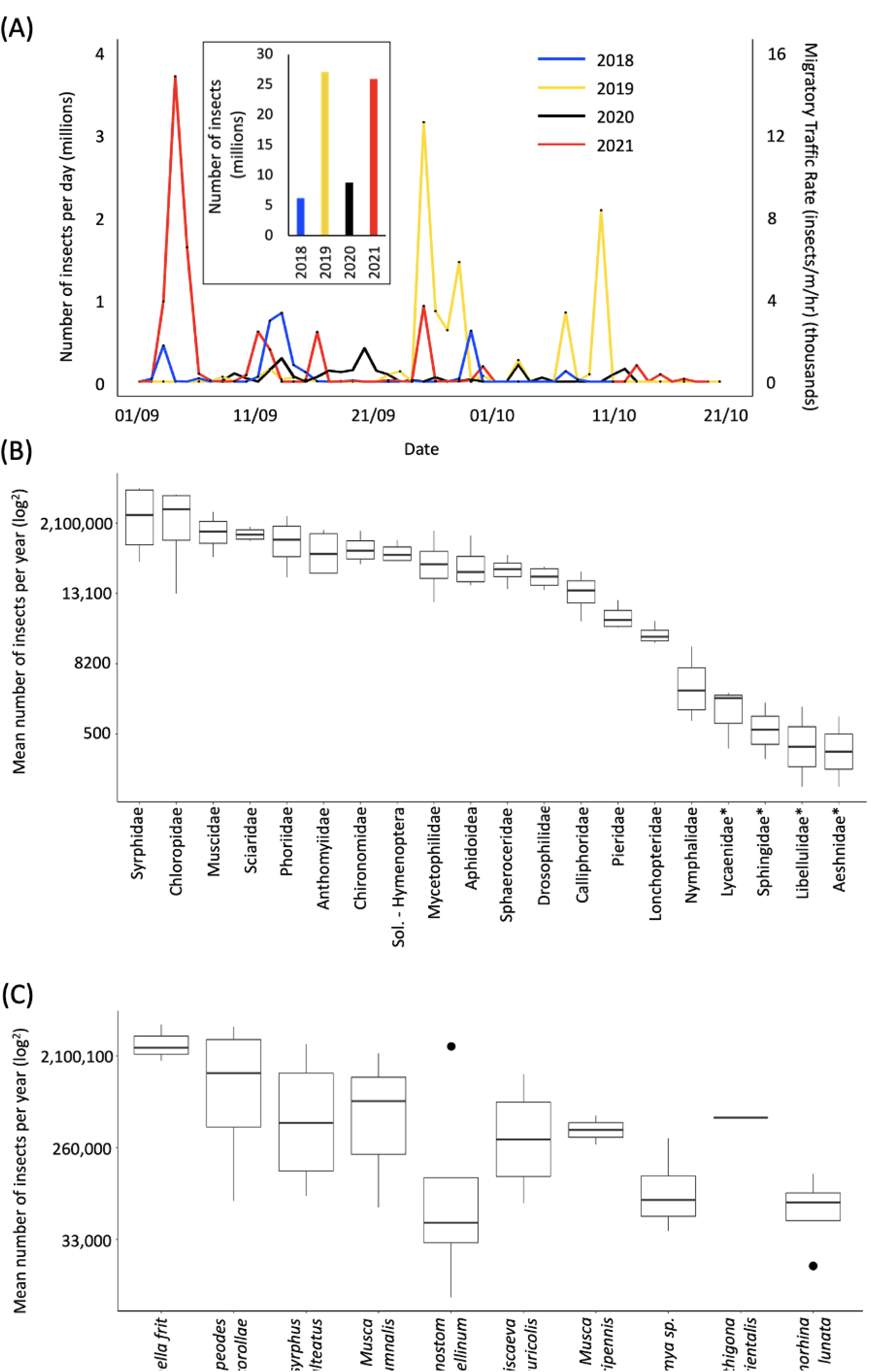
Migratory insect bioflows. (A) Number of insects and migratory traffic rates per day across four years based on video trap data (not including the estimated missed insects or butterflies). (B & C) Box and whisker plots detailing the variation in insect families (B) and the top ten most numerous species (C) across each season.

Of the migrant Diptera, Syrphidae were the most abundant, averaging 3.1 million per year and making up 20% of the migratory assemblage (Figure 3B). Chloropidae, Sciaridae, Muscidae and Phoriidae all averaged over a million individuals each year, with lower numbers seen for the Anthomyiidae, Mycetophilidae, Drosophilidae and Calliphoridae. Of the butterflies, the most abundant family were the Pieridae (*Colias croceus, Pieris rapae* and *P. brassicae*) at 55,000 per year while the libellulid *Sympetrum striolatum* (common darter dragonfly) was the most abundant dragonfly at 400 per year. At the species level, the most abundant in the bidirectional malaise traps were the *Oscinella* spp. grass flies (Chloropidae) and the hoverflies *Eupeodes corollae* and *Episyrphus balteatus* (Syrphidae), while flies from the Muscidae (such as *Musca autumnalis*) and from the Anthomyiidae (such as *Pegomya* sp.) were also abundant (Figure 3C). Larger dipteran migrants (*Eristalis tenax* and *Scaeva* spp.) were, in general, able to avoid the malaise trap despite being present in substantial numbers in the video traps. In general, sex ratios were skewed toward females for *Melanostoma mellinum* (F 65%, M 35%), *Eupeodes corollae* (F 95%, M 5%) and *Musca autumnalis* (F 72%, M 28%). In total, all migratory taxa comprised an average of 140 kg of biomass each year.

### The effect of environmental factors on the number of insects migrating

To understand the environmental factors important for migration we investigated the influence of average daily temperature, total daily sunshine, average wind speed and daily precipitation. We analysed the relationship of these factors with numbers of migrating insects (both camera trap and butterfly counts combined). Each of the explanatory variables were found to significantly affect the number of insects migrating through the pass (p = <0.0005) (Figure 4A-D). Comparison of the Aikaike’s Information Criterion (AIC) identified temperature as the variable that most affects the count number (p = <0.001, AIC: 71,808,928 (when temperature is removed) vs. all in: AIC: 60,027,687) showing a strong positive relationship (Figure 4A). This is followed by a positive correlation with sunshine and negative correlations with windspeed, and precipitation (p = <0.001, AIC: 67,554,952; p = <0.001, AIC: 61,995,988; p = <0.001, AIC: 61,236,263 respectively, Figure 4B-D).

**Figure 4.**
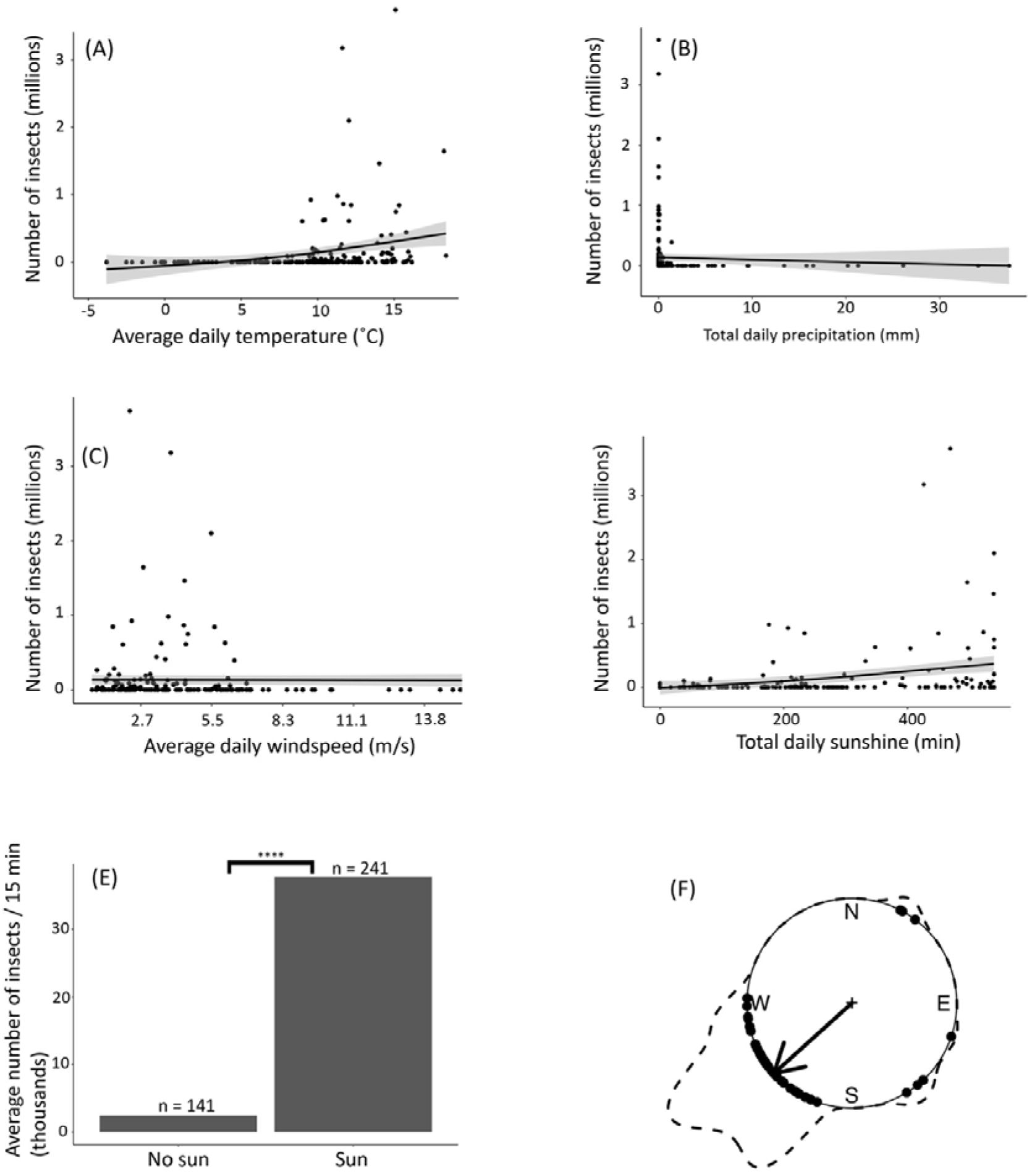
The relationship between the number of insects migrating through the Pass of Bujaruelo and meteorological variables. (A) average daily temperature, (B) total daily precipitation, (C) average daily windspeed, and (D) total daily sunshine. Shaded areas represent 95% confidence intervals, and the results are based on a generalised linear model with a Poisson distribution. (E) The average number of insects (per 15 minutes) when sunlight was present in the pass and when the sunlight was absent. (F) The mean wind heading for each mass migration day (black dots) was correlated with the total daily insect count (dotted line). The circular plot shows mean wind origin on ‘mass-migration’ days (arrow).

**Figure 5.**
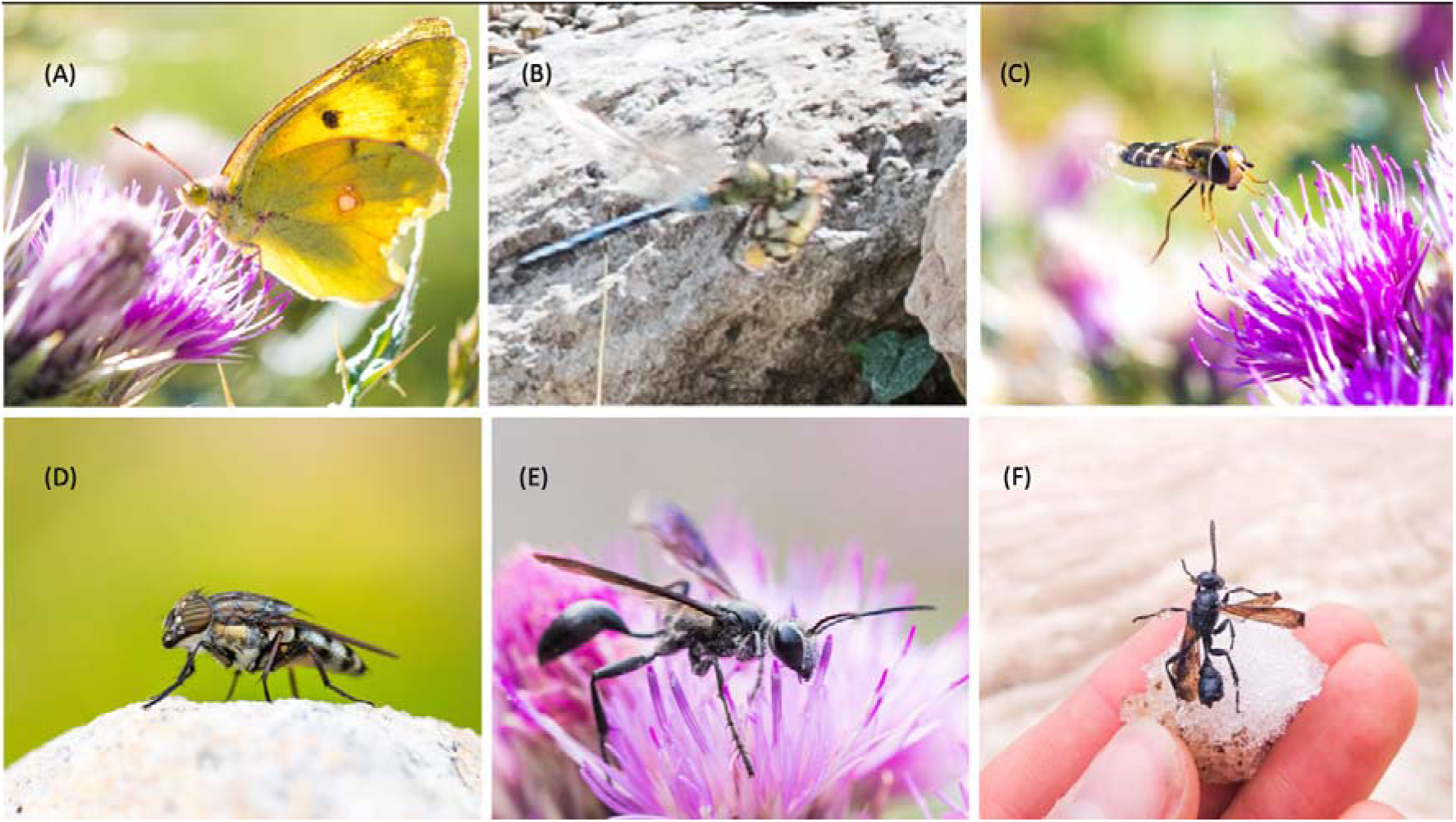
Migratory insects of the Pyrenees. (A) Clouded yellow (Colias croceus), (B) Migrant hawker (Aeshna mixta) eating a clouded yellow (Colias croceus), (C) Pied hoverfly (Scaeva pyrastri), (D) Locust blowfly (Stomorhina lunata), (E) Isodontia mexicana feeding on Pyrenean thistle (Carduus carlinoides) within the Puerto de Bujaruelo pass. (F) Isodontia mexicana found dead on a glacier at ∼2750m above sea level.

To investigate the influence of wind direction we analysed the relationship with ‘mass migration’ versus ‘non-mass migration’ events (see Methods). In total, 38 days were classed as mass migration events and 162 as non-mass migration events (Figure 4F). The majority of insects were found to be moving through the pass when a headwind (from the SW) was present (Figure 4E black arrow and dots; Rayleigh test: 227.9°, r = 0.7889, p = <0.001). On non-mass migration days, wind directions were random and not statistically significant (Rayleigh test: r = 0.134, p = >0.05) with peak migration rates 43.5-fold smaller than on mass migration days. On seven occasions a tail wind was present, and insects were recorded, but on all these occasions the windspeed was light, registering under 3.3 m/s (Figure 4F). To investigate the role of wind over a broader scale we visualised wind patterns using the Web application ‘<u>Ventusky</u>’ over Western Europe, centred on the Pyrenees. On 31 out of 38 days mass migration days, there was a tailwind or low windspeeds local to the northern (French) side of the mountains which met a headwind coming up from the south (Spain) along the ridgeline of peaks. This suggests that, on these occasions, the insects could utilise the favourable tailwind to reach the tops of the mountains before meeting headwinds at higher altitudes (Figure S2).

To account for fine scale variation in the effect of direct sunshine on migrant numbers we utilised 15-minute counting periods from the migration camera trap data. We found that the amount of sunshine significantly affected the numbers of insects moving through the pass (Figure 4E). Each 15-minute counting period when it was sunny contained 16-fold more insects on average than when the sun was hidden (t = -6.8, p = <0.0005) (Figure 4E).

Given the strong influence of temperature on migratory activity we tested if this factor, along with other meteorological conditions, might explain the 4.4-fold annual variation in migrant numbers. No variables were found to be significant given the current data, however a non-significant trend between Northwest European average August and September temperatures and the number of insects moving through the pass was identified (Figure 4G). To further investigate this, and as the majority of the migrants identified were hoverflies, we used a long term (10 year) entomological radar dataset consisting of detections of migratory hoverflies over the UK taken from (5). In this data set we uncovered a significant positive correlation between the abundance of migratory hoverflies and temperature in the autumn (R_38_= 0.454; *p* =<0.005) (Figure 4H).

### Estimating the number of insects crossing the entire Pyrenean Mountain range

Next, we used these variables to estimate the total numbers of insects crossing the whole Pyrenean range. Our calculations suggest that, between September and October, in all years there were an average of 8.75 days of headwinds when the conditions were suitable for migration where an average of 6.1 billion insects traversed the Pyrenees and an average of 12.5 days of suitable tailwinds where an average of 8.5 billion insects traversed the Pyrenees. In total the mean annual number of insects crossing the entirety of the Pyrenees in the autumn was estimated at 14.6 billion, equating to 125 tons of biomass. The year-by-year breakdown is displayed in Supplementary table S7.

## Discussion

More than 70 years after it was first discovered as an important insect migration hotspot (15) we returned to the Pass of Bujaruelo to systematically quantify the flow of day-flying insects migrating through the pass and to gain an understanding of the scale of the entire diurnal migration along the Western European insect flyway. We estimate that during our observation period of September to October 2018-2021, an average of 17.1 million insects traversed the pass each year heading south into Spain, and presumably beyond (23), on their migratory journeys. In the following sections we discuss meteorological conditions that predict migration events in the mountains and the drivers of annual variation in abundance. We also discuss the diverse range of species involved, the scale of the Western European flyway and the ecological impact of the estimated 14.6 billion individual day-flying insects crossing the entire Pyrenean Mountain range each autumn.

### Comparisons to historic observations at mountain passes

A lack of systematic data collection performed in the Pass of Bujaruelo in the 1950s makes comparisons challenging, particularly for Diptera, as migration rates through the pass were only compared to other migrants: e.g., “at least twenty times, and perhaps a hundred times, as numerous as the dragonflies”, who themselves were moving at a rate of “at least several thousand an hour” (15). In contrast, studies focussing on hoverflies in the 1960s and 70s in the Alps utilised a similar, but larger, intercept trap to ours (2 m^2^ vs. 16 m^2^) (24), making comparisons more straight forward following eight times scaling of our numbers. The most common hoverfly in our study, *E. corollae*, with an average (over the 4 years) scaled count of 6100 is just over 46% of the average 13,228 individuals caught in the 12 years between 1962 and 1973 (8). In contrast, the most common hoverfly in the Alps study was *E. balteatus* with our counts reaching only 2.6% of the 107,602 individuals caught by Aubert (8). Comparing numbers from many years apart and from different sites is problematic and we do not know how contemporary numbers would compare across sites, though we suspect they might be more similar given the potential declines in the numbers of migratory Syrphidae since the 1970s (16). If these declines have been present consistently across Europe, the numbers moving through the Pass of Bujaruelo during the study period of the early papers (1950s) may have been many times more than what is currently observed. Of the other insect groups, we occasionally saw butterflies passing through in numbers exceeding the rate of 500/hr noted by Lack & Lack (15), but while huge numbers of dragonflies were twice mentioned in the 1950s papers, for example “an uncountable stream of [*Sympetrum striolatum* dragonflies]” (17), during our four years at the Pass of Bujaruelo, we rarely saw dragonflies passing in such numbers.

We found strong evidence of migratory behaviour in small species not noted during the 1950s’ studies, likely due to a lack of systematic sampling techniques. The Chloropidae, Phoriidae, Drosophilidae, Mycetophilidae and the Sciaridae all showed migratory behaviour while the Sphaeroceridae, Chironomidae, Lonchopteridae, solitary Hymenoptera, and the Aphidoidea all showed weak migratory behaviour. Previous studies have regarded these insects as being simply at the mercy of air currents without the ability to choose their own direction (11). However, the consistent appearance of these insects in the southward heading side of our bidirectional malaise trap despite headwinds, suggests that these smaller insects are capable of directed migration. Further studies are needed to ascertain how these smaller species can select favourable conditions for movement, the distances they may be travelling and if they utilise similar compasses to larger Dipteran migrants (25). Finally, on a few occasions, migration of interesting species was noted but as they occurred in small numbers, they fall outside of our criteria and records of these species were made on an ad hoc basis. Details of these migrants can be found in the Supplementary Results & Discussion.

### The Western European migratory assemblage and ecological roles

Pyrenean migration hotspots have been relatively poorly studied for insects despite the early recognition of these flyways (13,15). We identified 20 families of day-flying insects from five Orders showing migratory behaviour as judged by directional movement through the Pass of Bujaruelo. The ecological roles were dominated by pollinators that made up nearly 90% of the insects recorded followed by pest species (36%), decomposers (34%) and controllers of pests (20%) while all contributed to the transport of nutrients. A full discussion of these roles is presented in the Supplementary Results & Discussion. The spread of these ecological roles over large geographic areas, and through time, is likely to have major consequences for the functional of ecosystems. For example, migratory pollinators, can link geographically isolated plant populations through their movements, allowing for gene flow between these populations (26) while the movement of pest species may lead to direct damage to crops while also increasing rates of geneflow, increasing the chance of resistance arising and spreading (27). In addition, the movement of large amounts of biomass through the pass, estimated at a little over 140 kg and equating to 14 kg of Nitrogen and 1.4 kg of Phosphorous, forms part of a larger movement of insects that together represents the largest terrestrial bioflow (2), and has major potential benefits, including revitalising various ecosystems along with the arrival of migratory insects.

### Environmental factors influencing migratory intensity and total numbers

Environmental factors have been shown previously to affect the numbers of insects arriving at migratory hotspots (14). Our model from the Pyrenees showed that average daily temperature, total daily sunlight, average daily windspeed, and total daily precipitation, were significant in affecting the number of insects migrating through the pass. Further analysis of meteorological conditions revealed that temperature was the variable explaining the most variation in insect numbers, followed by sunlight, windspeed, and finally precipitation. Temperature was found to be positively correlated with insect number, a finding that is consistent with findings of migrating dragonflies in Latvia, hoverflies in North America, and with migratory insects of all taxa in Cyprus (6,14,28). The total daily sunlight’s positive correlation with insect number is also consistent with previous Pyrenean observations by Williams et al. (13) who stated that “sunshine is a major factor determining […] migration”, and findings that the effect of sunlight and temperature on insect migration is tightly linked (6,29). When analysing the effect of sunlight on insect numbers at a fine scale, we found that during each 15-minute counting period throughout the day when sunlight was present, insects were 16-fold more abundant than when there was no sun. Insects are often highly sensitive to changes in temperature due to the presence of sunlight, their small body size is particularly susceptible to heat loss, and in addition the sun may be required to orientate (25,30,31).

The topography of the Pyrenees means that on days when a tailwind is present, the insects can ride updrafts over the >3000 m peaks and we observed migration of flies and butterflies through the Brèche de Roland (2,804 m) and over the peak of Taillon (3,144 m) on multiple occasions. On headwind days, however, insects are forced down low into their ‘flight boundary layer’ (32) to avoid the winds and are guided by topographical features through mountain passes like Bujaruelo. Therefore, in contrast to studies in Latvia and Cyprus (amongst others), insects were recorded almost exclusively during headwinds (Figure 4A) (6,14). Insects are well known to choose favourable tailwinds to aid their migratory journeys (2,33–37), but our observations show that, in the mountains and under certain circumstances, this is not always the case. The most likely explanation is that of orographic wind, the process by which a mass of air is lifted by a large geographical feature such as a mountain range, creating winds that blow towards the mountain tops (38). These local wind conditions are therefore as important (if not more so) than the synoptic-scale tailwinds blowing from France into Spain, with regards to the insects migrating through the Pyrenees. Therefore, while insects can utilise a tailwind to help power their flight south, on reaching the ridgeline they may need to counteract a headwind to continue their journey.

The early papers noted large-scale insect migration through the Pass of Bujaruelo exclusively during a headwind and that the insects flew within the flight boundary layer where the windspeed was weaker, suggesting windspeed is an important factor for the insect migrants (13,15). In the present study, windspeed was found to negatively correlate with the numbers of insects migrating, we believe this is a result of the insects usually needing to migrate into a headwind when moving through the pass. Studies on dragonflies in USA and hoverflies over the UK during the autumn found that they too prefer to migrate in lower windspeeds, potentially allowing the insects to have more control on their flight direction (36,39). Finally, total daily precipitation was found to negatively correlate with the numbers of insects migrating through the pass. Variation in numbers over the whole season for the four years ranged 4.4-fold and loosely mirrored August and September temperatures from a higher latitude. This trend was strongly supported by analysis of radar data showing a significant positive correlation between numbers of hoverflies migrating above Northwest Europe in autumn and higher temperatures in August and September from the same area. In summary, predictors of large migration events are warmer temperatures in Northwest Europe and, on the pass, sunny conditions, a light headwind from the south of less than 7 m/s at 2m elevation above ground, no rain and a temperature in the valley of >13°C. This predictive model can be applied to other passes in high mountain ranges (Figure S3) and, in the next section, we use it to predict total insect numbers crossing the Pyrenees.

### Total numbers traversing the Pyrenees and the Western European Insect Flyway

Numerous studies have found a consistent south or southwest bias in migratory insect headings across Europe (37,40–42) suggesting insects from a large geographical area are filtered down into the Iberian Peninsula, passing through the Pyrenees each year. This makes insect migration hotspots within the Pyrenees important locations for censusing species and monitoring numbers. We estimate that the total number of insects moving across the Pyrenean Mountain range reaches an average of 14.6 billion each year. This number is of comparable size to those from radar studies across a similar sized area. Hu et al. (2) found that ∼100 tons of both diurnal and nocturnal migratory insects leave southern England each year for the continent. Our estimates of the total biomass of diurnal migratory insects moving across the Pyrenees each autumn equates to around 125 tons, and this number does not include nocturnal migrants. It is unlikely that nearly all the insects moving across the Pyrenees are of British origin, so we are surely missing a great deal based on our Pyrenean predictions. Reasons for this may include (1) high mortality on the journey; or (2) that the majority of the insect migrants move along the eastern and western edges of the Pyrenees where the altitude is lower. In fact, Williams et al. (13) observed some southward movement of butterflies in October along the coast at Argeles-sur-Mer where the Pyrenees descend down to the Mediterranean Sea. Similar southward movements occur along the Atlantic seaboard (43,44). Also, there may well be more migration through lower, wider Pyrenean passes, but it is less easy to monitor insect migration within these. Additionally, our estimates of numbers are based on weather conditions recorded from the Pass of Bujaruelo, a high-altitude pass in the centre of the Pyrenees. As it is likely that weather conditions will be more favourable to insect migration at lower altitudes, our predictions are likely underestimating the number of insects moving. However, to fully research this, extensive deployment of monitoring resources and techniques are needed (45).

## Conclusion

Our four-year study of insects migrating through a Pyrenean Mountain pass has revealed a remarkable diversity and abundance of migratory insect taxa. Additionally, based on the south-southwest bias of insect movement during the autumn season in Europe, the insects migrating through the Pyrenees are likely to have originated from across Western Europe (36,40). More long-term studies of migratory hotspots are required to expand the monitoring of insect populations and to avoid ‘shifting baseline syndrome’ (46). We suggest studies incorporate use of low-cost migration cameras to collect data during headwinds and, where possible, entomological radar to fully account for the migration in tailwind conditions, though we recognise the potential difficulty in locating and maintaining such technology in remote locations. Our study has shown that many more insect taxa show directed migratory behaviour than we were previously aware of and that research into the ecological benefits of these insect migrants is of importance, especially to the agricultural and horticultural sectors. These findings highlight the benefits accrued from insect migration (including nutrient transfer, pest control, pollination and decomposition) and provides key predictors of migratory events, vital in understanding population changes the era of anthropogenically induced environmental change.

## Acknowledgements

We would like to thank the Parc national des Pyrénées and the Gobierno de Aragon for allowing us to perform this work in their beautiful national parks. Additionally, we thank Gao Hu for supplying radar data used within this study. WLH would like to thank Sarah Hawkes for continuous support and insightful comments.

## Funding

This work was supported through grants to KRW from the Royal Society University Research Fellowship scheme (grant no. UF150126). TD and WLSH were supported by awards to KRW from the Royal Society: a Fellows Enhancement Award (RGF\EA\180083) and a Research Grant for Research Fellows (RGF\R1\180047), respectively. RM was supported through the NERC GW4+ Doctoral Training Partnership. Rothamsted Research receives grant-aided support from the UK Biotechnology and Biological Sciences Research Council.

## Notes

### Competing Interest Statement

The authors have declared no competing interest.

