## Supplementary file for "The most remarkable migrants – systematic analysis of the Western European insect flyway at a Pyrenean mountain pass"

**Supplementary Methods**

**Determining migrants from the bidirectional malaise trap**

To be included in the analysis, insect taxonomic groups had to be recorded >100 times in the bidirectional malaise trap counts. Migrants were determined based on a ‘southward score’ which is the percentage caught heading south in a wind category (headwind or tailwind) minus the percentage caught heading north in the same wind category.

$$Headwind southward score=Percentage heading south-Percentage heading north$$

$$Tailwind southward score =Percentage heading south-Percentage heading north$$

Taxonomic groups were then sorted into four categories: ‘strongly migratory’, ‘migratory’, ‘weakly migratory’, and ‘non-migratory’ based on the southward scores under different wind conditions. ‘Strongly migratory’ includes taxonomic groups with a southward score of ≥90 in headwind conditions but with <100 individuals caught in tailwinds presumably as they can utilise these favourable tailwinds to fly over the highest peaks and so do not use the pass. ‘Migratory’ includes taxonomic groups with a southward score between 50 – 89 in headwind conditions and a score of ≥50 in tailwind conditions indicating that they still use the pass in tailwinds. ‘Weakly migratory’ includes taxonomic groups with a southward score of <50 in a headwind condition and ≥70 in tailwind conditions. ‘Non-migratory’ is classified as <50 in a headwind condition and <70 in tailwind conditions.

**Quantifying numbers of insects annually traversing the Pyrenees**

To quantify the numbers of insects passing over the entirety of the Pyrenees each autumn we identified the location of the valley mouth (located at: 43°04'46.9"N 0°02'43.4"W) where the migratory assemblage moving through the Pass of Bujaruelo would first begin to be channelled up the valley towards the pass. This valley mouth is 1.2 km wide. We make the following assumptions: (1) As the insects are channelled by the steep-sided valley towards the Pass of Bujaruelo, the number of insects entering the valley at the mouth (1.2 km) is equal to that moving through the 30m wide pass, and; (2) that the number of insects recorded migrating is at a constant rate across the Pyrenees, a presumption likely to lead to an under-estimation, as the edges of the Pyrenees have less topological barriers to restrict migration and as weather conditions in the high mountains are frequently unsuitable for migration. Based on these assumptions we calculate the flow over the 430 km length of the Pyrenees on headwind days based on the numbers at the pass, that is as the flow through (430 km / 1.2 km) 358, 1.2 km segments.

$$Insects moving through Pyrenees on headwind days=yearly average x 358$$

To calculate the number of insects on tailwind days we assume that the number of insects migrating on a tailwind day equals the number on headwind days per 1.2 km segment. However, for migration to occur on a headwind day various meteorological criteria must be met based on our measurements (see Figure 7): the wind must be from between 290˚- 60˚, rainfall must be below 1.4 mm per day and the temperature at the mouth of the valley must be higher than 10˚C. This left 11 viable days in 2018, 14 days in 2019, 11 days in 2020 and 14 days in 2021.

$$Insects moving through Pyrenees on tailwind days = Daily average numbers of insects that year x number of viable days x 358$$

Finally, we calculate the annual movement of insects through the Pyrenees.

$$Total anual number of insects moving through whole Pyrenees= the total number of insects (headwind) + total number of insects (tailwind).$$

**Supplementary Tables**

**Table S1**. Insect taxa from the Pyrenees whose numbers caught in the malaise trap did not exceed 100 individuals and so were not included in the migration analysis.

| Insect taxa | Average number of insects per season | Percentage of total assemblage |
| --- | --- | --- |
| Coleoptera: Staphylinidae | 26.5 | 0.3 |
| Diptera: Bibionidae | 24.5 | 0.28 |
| Homoptera (Leafhoppers) | 22.5 | 0.26 |
| Diptera: Dryzomyidae | 21.75 | 0.25 |
| Diptera: Tipulidae | 19.5 | 0.22 |
| Hymenoptera: Formicidae | 18.75 | 0.21 |
| Diptera: Fanniadae | 18.3 | 0.21 |
| Diptera: Cecidomyiidae | 16.5 | 0.19 |
| Diptera: Platystomatidae | 13 | 0.15 |
| Diptera: Scathophagidae | 8.5 | 0.09 |
| Diptera: Pallopteridae | 8 | 0.09 |
| Diptera: Sepsidae | 7 | 0.08 |
| Diptera: Tephritidae | 6.2 | 0.07 |
| Hymenoptera: Argidae | 5.7 | 0.06 |
| Hymenoptera: Apidae | 4.5 | 0.05 |
| Arachnida: Opilionidae | 3.5 | 0.04 |
| Diptera: Psychodidae | 3.5 | 0.04 |
| Hymenoptera: Symphyta (non-Argidae) | 3.25 | 0.03 |
| Diptera: Empidae | 3 | 0.03 |
| Arachnida: Acari | 2.7 | 0.03 |
| Diptera: Tachinidae | 2.2 | 0.02 |
| Diptera: Dolichopodidae | 2 | 0.02 |
| Diptera: Acroceridae | 2 | 0.02 |
| Diptera: Psilidae | 1.7 | 0.02 |
| Hymenoptera: Mymaridae | 1.5 | 0.01 |
| Diptera: Hippoboscidae | 1 | 0.01 |
| Lepidoptera: Crambidae | 1 | 0.01 |
| Lepidoptera: Pterophoridae | 1 | 0.01 |
| Coleoptera: Coccinelidae | 0.7 | 0.008 |
| Arachnida: Thomisidae | 0.7 | 0.008 |
| Hymenoptera: Vespidae | 0.6 | 0.007 |
| Diptera: Stratiomyidae | 0.5 | 0.005 |
| Arachnida: Gnaphosidae | 0.5 | 0.005 |
| Hymenoptera: Sphecidae | 0.5 | 0.005 |
| Hymenoptera: Crabronidae | 0.5 | 0.005 |
| Neuroptera: Ascalaphidae | 0.5 | 0.005 |
| Lepidoptera: Erebidae | 0.3 | 0.003 |
| Coleoptera: Curculionidae (Apionidae) | 0.25 | 0.002 |
| Diptera: Ephrydidae | 0.25 | 0.002 |
| Diptera: Tabanidae | 0.25 | 0.002 |
| Dermaptera: Forficulidae | 0.25 | 0.002 |
| Diptera: Conopidae | 0.25 | 0.002 |

**Table S2.** Insect types with 'strong migratory behaviour’. Average numbers are given based on scaled video trap counts including the ‘missed insects’ (see main paper) or, where indicated with an *, on butterfly counts.

| Insect group/species | Dominant species | Average number per season | Percentage of total migratory assemblage | Southward score (Headwind:Tailwind) |
| --- | --- | --- | --- | --- |
| Diptera: Syrphidae | *Eupeodes corollae* | 3.1 million | 20.2 | 100:(n/a) |
| Lepidoptera: *Vanessa atalanta** | *Vanessa atalanta* | 5960 | 0.04 | 100:(n/a) |
| Lepidoptera: Lycaenidae* | *Lampides boeticus* | 5024 | 0.001 | 100:(n/a) |
| Diptera: Calliphoridae | *Stomorhina lunata* | 0.14 million | 0.9 | 98:(n/a) |

**Table S3.** Insect groups with ‘migratory behaviour’. Average numbers are given based on scaled video trap counts including the missed insects or, where indicated with an *, on butterfly counts. n/a indicate that less than 100 individuals were recorded.

| Insect group/species | Dominant species | Average number per season | Percentage of total migratory assemblage | Southward score (Headwind:Tailwind) |
| --- | --- | --- | --- | --- |
| Diptera: Chloropidae | *Oscinella frit* | 2.7 million | 17.4 | 89:97 |
| Diptera: Sciaridae | *Sciara sp.* | 2 million | 13.2 | 54:85 |
| Diptera: Muscidae | *Musca autumnalis* | 1.5 million | 9.4 | 97:74 |
| Diptera: Phoriidae | Unidentified | 1.1 million | 7 | 90:81 |
| Diptera: Anthomyiidae | *Delia spp.* | 0.7 million | 4.6 | 94:56 |
| Diptera: Mycetophilidae | Mycetophilinae | 0.5 million | 3.5 | 71:94 |
| Diptera: Drosophilidae | *Scaptodrosophila spp.* | 0.25 million | 1.7 | 89:98 |
| Lepidoptera: *Colias croceus** | *Colias croceus* | 0.04 million | 0.3 | 100:100 |
| Lepidoptera: *Pieris sp.** | *Pieris rapae* | 15,000 | 0.1 | 100:100 |
| Lepidoptera: Sphingidae* | *Macroglossum stellatarum* | 2560 | 0.004 | (n/a):(n/a) |
| Odonata: Libellulidae* | *Sympetrum striolatum* | 392 | 0.001 | (n/a):(n/a) |
| Odonata: Aeshnidae* | *Aeshna mixta* | 272 | 0.002 | (n/a):(n/a) |

**Table S4.** Insect types with ‘weak migratory behaviour’.

| Insect taxa |  | Percentage of total assemblage | Southward score (Headwind: Tailwind) |
| --- | --- | --- | --- |
| Diptera: Sphaeroceridae | 0.38 million | 2.2 | 36:96 |
| Diptera: Chironomidae | 1.1 million | 6.6 | 38:74 |
| Solitary Hymenoptera | 0.88 million | 5.1 | 32:72 |
| Diptera: Lonchopteridae | 0.1 million | 0.6 | (n/a):96 |
| Hemiptera: Aphidoidea | 0.7 million | 4.2 | -17:84 |

**Table S5.** Breakdown of the ecological roles performed by the migrant insects moving through the Pass of Bujaruelo.

| **Role** | **Major group** | **% of total** | **Average number migrating through the pass per season in millions (Total)** |
| --- | --- | --- | --- |
| Pollinator | Diptera: Syrphidae | 19.9 | 7.3 (29.2) |
|  | Diptera: Chloropidae | 17.1 | 6.3 (25.1) |
|  | Diptera: Sciaridae | 13 | 4.8 (19.1) |
|  | Diptera: Muscidae | 9.3 | 3.4 (13.7) |
|  | **Total** (including minor groups) | **87** | **14.8 (59 million)** |
| Pests | Diptera: Sciaridae | 13 | 4.8 (19.1) |
|  | Hemiptera: Aphidoidea | 4.5 | 1.7 (6.6) |
|  | Diptera: Chloropidae | 17.1 | 6.3 (25.1) |
|  | Lepidoptera: Pieridae | 1.1 | 0.4 (1.6) |
|  | **Total** | **36** | **6.1 (24.5 million)** |
| Decomposers | Diptera: Sciaridae | 13 | 4.8 (19.1) |
|  | Diptera: Muscidae | 9.3 | 3.4 (13.7) |
|  | Diptera: Chironomidae | 7.2 | 2.7 (10.6) |
|  | **Total** (including minor groups) | **33.6** | **5.5 (22.8 million)** |
| Pest predators | Diptera: Syrphidae (*Eupeodes corrolae, Episyrphus balteatus, Sphaerophoria scripta*) | 15.6 | 5.7 (22.9) |
|  | Solitary Hymenoptera | 5.6 | 2.1 (8.2) |
|  | **Total** (including minor groups) | **22.2** | **3.8 (15.1 million)** |

**Table S6.** Breakdown of the number of insects per season moving through the Pass of Bujaruelo.

| **Year** | **Insect count (Missed insects) (millions)** |
| --- | --- |
| 2018 | 2.3 (6.2) |
| 2019 | 10.1 (27.1) |
| 2020 | 3.3 (8.8) |
| 2021 | 9.7 (25.9) |
| **Average** | 6.35 (17.1) |
| **Total** | 25.4 (68) |

**Table S7.** The annual variation of insects traversing the whole Pyrenean Range.

| **Year** | **Number of suitable headwind migration days (number of insects, (Missed Insects (M.I.)) in billions)** | **Number of suitable tailwind migration days (number of insects (Missed Insects (M.I.)) in billions)** | **Total numbers of insects traversing the Pyrenees (including Missed Insects (M.I.) in billions)** |
| --- | --- | --- | --- |
| 2018 | 8 (0.8 (2.2)) | 11 (1.1 (3)) | 1.9 (5.3) |
| 2019 | 10 (3.6 (9.7)) | 14 (5 (13.5)) | 8.7 (23.3) |
| 2020 | 6 (1.2 (3.1)) | 11 (2.1 (5.7)) | 3.3 (8.9) |
| 2021 | 11 (3.5 (9.3)) | 14 (4.4 (11.8)) | 7.9 (21.1) |
| **Average** | 8.75 (2.3 (6.1)) | 12.5 (3.2 (8.5)) | 5.5 (14.6) |
| **Total** | 35 (9.1 (24.3)) | 50 (12.8 (34.2)) | 21.9 (58.5) |

**Supplementary Figures**

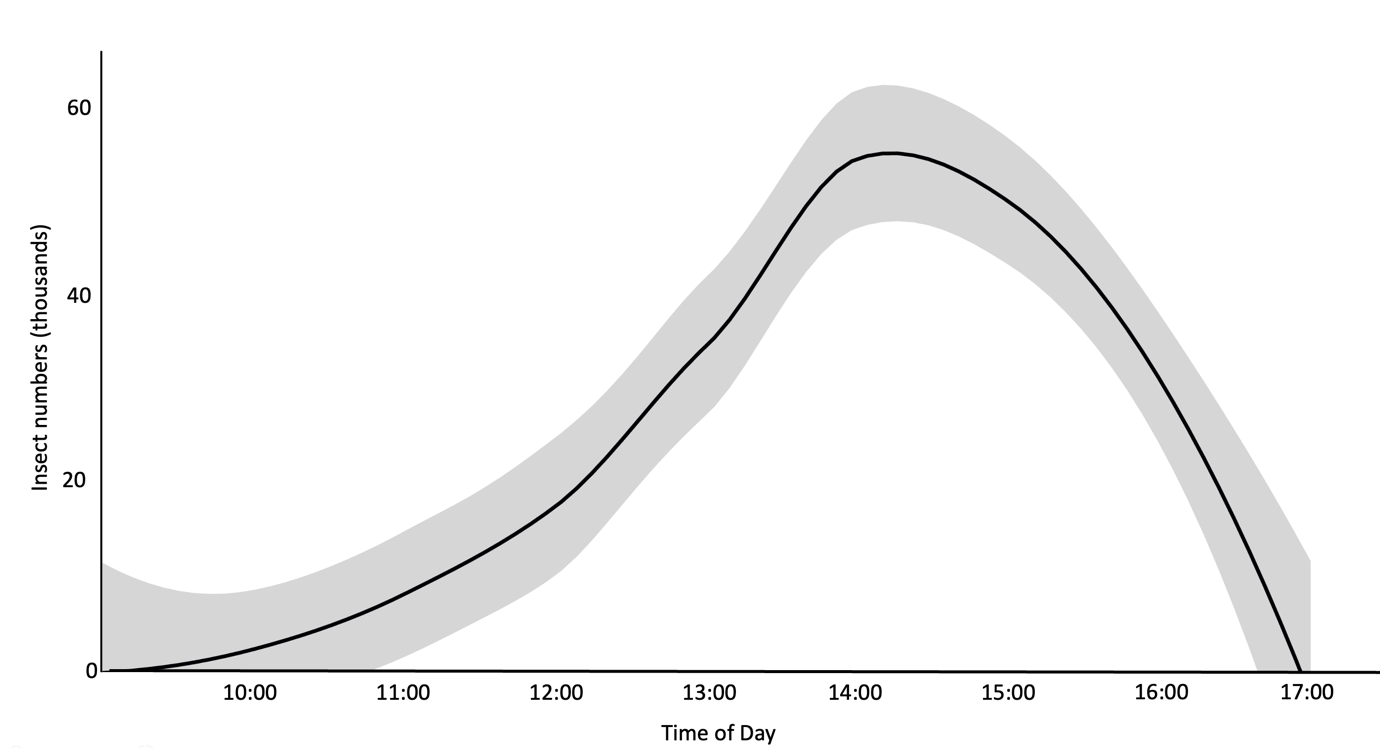

**Figure S1**. The temporal distribution of insects throughout the 09:00 to 17:00 period based on ‘mass migration’ events across four years of data collection.

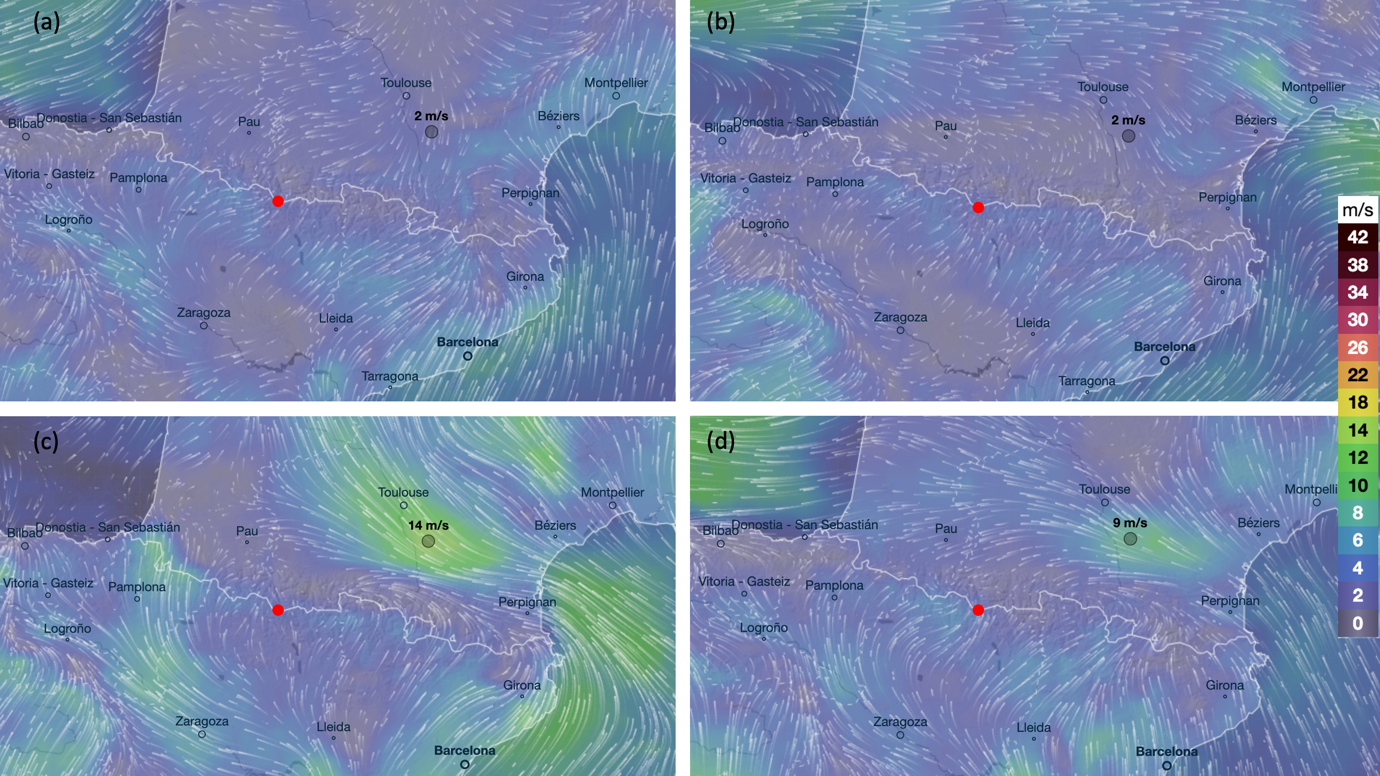

**Figure S2. Visualisation of wind patterns 10 metres above the ground on the four largest mass migration days.** To investigate the role of wind over a broader scale we visualised wind patterns using the Web application ‘[Ventusky](https://www.ventusky.com/)’ over Western Europe, centred on the Pyrenees on 38 days of mass migration events through the Pass of Bujaruelo. On 31 out of these 38 days, there was a tailwind or low windspeeds local to the northern (French) side of the mountains which met a headwind coming up from the south (Spain) along the ridgeline of peaks. This suggests that, on these occasions, the insects could utilise the favourable tailwind to reach the tops of the mountains before meeting headwinds at higher altitudes.(A) 5 Sept 2021, (B) 26 Sept 2019, (C) 11 Oct 2019, (D) 6 Sept 2021. Colour gradients signify wind speeds in m/s. The red dot indicates location of the Pass of Bujaruelo.

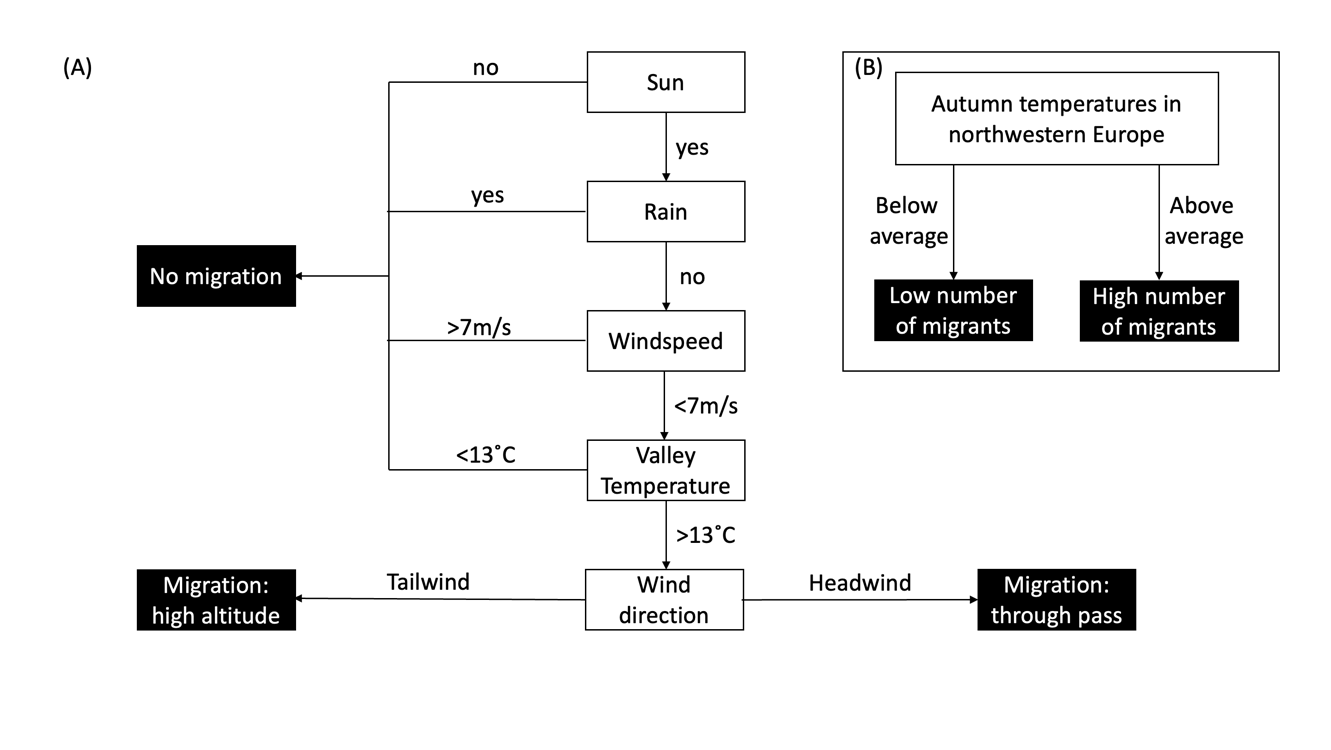

Figure S3. **A flow chart to predict insect migration through the Pyrenees.** (A) A flow chart predicting insect migration on a daily basis. (B) A flow chart predicting number of migrants annually. On a broader geographic scale (i.e., across the Pyrenean Mountain range) when a tailwind meets a headwind on the Pass of Bujaruelo, mass migration of insects occurs.

**Supplementary Results & Discussion**

**Incidental insect movements**

On a few occasions, migration of interesting species was noted but as they occurred in small numbers, they fall outside of our main quantification criteria. Records of these species were made on an ad hoc basis: catching specimens to confirm identity and performing counts of individuals if large numbers were visible. The ichneumonid wasp *Obtusodonta equatoria* was found moving south through the Pass of Bujaruelo in relatively large numbers, particularly on the 9 October 2020 when they occurred at a rate of 3.2 per minute during a count of 5 minutes across a two metre transect, or scaled over the total Pass width, at a rate of 48 per minute. Directed movements southwest of the buff-tailed bumblebee (*Bombus terrestris*) were also often observed. Only queens of this species were seen and no *B. terrestris* nests were found in the immediate vicinity below the pass. Finally, individuals of the invasive grass carrying wasp *Isodontia mexicana* (Sphecidae) were also observed moving south through the pass. On one occasion, an individual was found dead on the glacier of the Brech de Rolando, ~2750 m above sea level. No vegetation or suitable feeding/breeding habitat was present at this elevation.

**The Western European migratory assemblage and ecological roles**

Pyrenean migration hotspots have been relatively poorly studied for insects despite the early recognition of these flyways (1,2). We identified 20 families of day-flying insects from five Orders showing migratory behaviour as judged by directional movement through the Pass of Bujaruelo. To understand the ecological impact of these movements we identified the ecological roles played by these families (Supplementary table S5) and we examine each within a migratory context in the following sections.

Nearly 90% of the insects recorded migrating through the Pass of Bujaruelo were pollinators. The major group was the hoverflies (Syrphidae), but with many others such as the butterflies and other Dipteran families such as the Muscidae also playing a role (3–5). Although low in numbers, the presence of highly efficient pollinators such as queen buff-tailed bumblebees (*Bombus terrestris*) were noted with directed flight southwest through the pass. This suggests that the queens are moving southwards for the winter, adding further evidence for bumblebee migration (6). Bumblebee migration has many implications including their exceptional pollination abilities and the continual replenishment of bumblebees to even the most intensively farmed areas from highly productive areas far away (6,7). Migratory pollinators, particularly species known to take part in long-distance movement such as the syrphids, may be particularly beneficial as they can link geographically isolated plant populations through their movements, allowing for gene flow between these populations (8). This gene flow can aid plant populations through reduction of inbreeding depression, general upkeep of population health, and they may have the potential to introduce new alleles to populations allowing them to adapt to the changing climate (e.g., alleles for drought resistance) (8,9).

Nearly 36% of the insects recorded migrating through the Pass of Bujaruelo can be classed as pest species, that is they are deemed detrimental to humans or human concerns. The major pest group migrating through the pass were the grass flies (Chloropidae – 17% of all migrants in pass), specifically *Oscinella frit* which consumes various cereals, grasses, and spring-sown maize as a larva (10,11). The aphids (Hemiptera) were also found to show weak migratory behaviour through the pass, suggesting some form of preferred directionality in their movements. Aphids are major pest species on a variety of crops and cause loses worldwide (12). Of the butterflies, the small white (*Pieris rapae*) is a known pest of Brassicaceae crops such as cabbages. The migration of these pest species is not only directly damaging to the crops themselves at their migratory destinations, but as a result of the migratory behaviour, the effectiveness of pesticides against these pests may be reduced (13). This is because migratory species often have large and diverse populations with a high rate of geneflow, allowing resistance to spread (13).

A little over one third of the insects recorded migrating through the Pass of Bujaruelo played an ecological role in decomposition. The major migratory decomposers found during the current study were the Sciaridae (13% of all migratory insects), and the Muscidae (9%), but also contained the Calliphoridae (2%) and Eristaline hoverflies such as *Eristalis tenax* (0.1%). The role of decomposition by migratory insects can be split into two groups: the decomposition of decaying vegetable matter and animal waste, and the decomposition of decaying animal flesh. In the former category the Sciaridae, Muscidae, and Eristaline hoverflies are important species. Sciaridae larvae are known to develop on a variety of terrestrial organic substances such as mammal faeces (14), and fungal or vegetable matter (15,16). The commonest Muscidae fly in our study was *Musca autumnalis* which lays its eggs in cattle dung, helping to decompose this abundant organic matter (17). Eristaline hoverfly larvae develop in decaying organic matter such as stagnant pools and livestock slurry pits (18). This breakdown of organic matter can reduce the impacts of eutrophication, acidification, and global warming through preventing methane release. In the decomposition of decaying animal flesh category, we documented blowflies such as *Calliphora spp.* and *Lucillia sp.*, who lay their eggs within carrion which their larvae then consume (19,20). This scavenging behaviour by the migrants leads to the redistribution of nutrients over large distances.

Over 20% of the insects recorded migrating through the Pass of Bujaruelo can be classed as natural enemies of pests. Primarily, these consist of the aphidophagous hoverflies such as *Eupeodes corollae, Episyrphus balteatus, Scaeva selenitica,* and *Scaeva* *pyrastri,* who together comprise nearly 14% of the total migratory insect assemblage. These hoverflies prey on aphids which feed upon crops. For example, *Eupeodes corollae* and *Episyrphus balteatus* migrating to southern England during the springtime, and the subsequent generations, have been shown to consume aphids equalling 20% of the early spring aphid population (21). Other migrants present in the pass with known pest controller behaviour include *Stomorhina lunata,* (Calliphoridae) whose larvae feed upon migratory locusts, the wasp *Obtusodonta equatoria* (Ichneumonidae) that may follow and lay her eggs on the caterpillars of the also migratory and destructive pest turnip moth (*Agrotis segetum*) (22).

All the insects moving through the Pass of Bujaruelo are transporting nutrients. The dry body of a migratory insect consists of 10 % Nitrogen and 1% Phosphorous, elements which are limiting to plant growth. Insects migrate in huge numbers as has been shown here, and in other studies which have recorded movements of trillions of individuals (23,24). As a result, these insects represent a rich source of nutrient input for ecosystems when, upon death, the nutrients of the insects are released into the ecosystem. The average weight of the total biomass moving through the Pass of Bujaruelo each year was a little over 140 kg, equating to 14 kg of Nitrogen and 1.4 kg of Phosphorous. The Syrphidae contributed the most to this total, with an average of 61 kg of biomass moving through the pass each year. This field is understudied but it is thought that the movement of migratory insects represents the largest terrestrial bioflow, equal to if not greater than those of the marine ecosystems (23), and has major potential benefits including revitalising high-latitude ecosystems with the arrival of migratory insects in the springtime.
